# FPGA Acceleration of GWAS Permutation Testing

**DOI:** 10.1101/2022.03.11.483235

**Authors:** Yaniv Swiel, Jean-Tristan Brandenburg, Mahtaab Hayat, Wenlong Carl Chen, Mitchell A. Cox, Scott Hazelhurst

## Abstract

**Motivation:** Genome-wide association studies (GWASs) analyse genetic variation over the genomes of many individuals in an attempt to identify single nucleotide polymorphisms (SNPs) associated with complex phenotypes. To capture a large amount of genetic variation and increase the chance of detecting associated SNPs, modern GWASs include millions of SNPs sampled from thousands of individuals. The cumulative probability of false associations increases with the number of SNPs included in the analysis. A GWAS, therefore, needs to control the number of false associations. Permutation testing is a straightforward and accurate method of controlling the false positive rate, but it is very computationally expensive (and slow) so there is a need for a permutation testing accelerator that can process modern GWAS datasets in reasonable time.

**Results:** FPGAs (Field-Programmable Gate Arrays) are reconfigurable integrated circuits which provide a high level of parallelisation that can be harnessed to accelerate GWAS permutation testing. This work presents an accessible FPGA-based tool (designed to run on a cloud-based AWS EC2 FPGA instance) that accelerates GWAS permutation testing for continuous phenotypes. The tool implements two known GWAS permutation testing algorithms: maxT permutation testing and adaptive permutation testing. The speed of the FPGA-based tool was compared to the speed of PLINK (a popular CPU-based tool) running on 40 Intel Xeon 4114 CPU cores using an imputed breast cancer dataset of 13.7 million SNPs sampled from 3652 individuals. For 1000 maxT permutations, the FPGA-based algorithm’s run time was 22 minutes while PLINK’s run time was almost 7 days; for 100 million adaptive permutations, the FPGA-based algorithm’s run time was 325 minutes and PLINK’s run time was about 8.5 days. For 700 million adaptive permutations of the same dataset (an almost unfeasible workload for PLINK running on a 40-core CPU) the run time of the FPGA-accelerated algorithm was 33 hours.

**Availability:** An EC2 AMI (FPGA_perm) in the us-east-1 region is available. Instructions, source code and sample data are available at https://github.com/witseie/fpgaperm.

## Introduction

Low cost, high density genotyping enables the collection of large datasets that provide a representative sample of the genetic variation across a population. A genome-wide association study (GWAS) attempts to associate single nucleotide polymorphisms (SNPs) with a phenotype using genome-wide data sampled from many individuals (1). Once a GWAS identifies phenotype-associated SNPs, the SNPs can be studied in more detail in order to gain insight into the genetic mechanisms underpinning complex phenotypes. GWASs have been widely applied to the association of SNPs with complex human diseases and other traits (2, 3). Identifying disease-associated variants can aid the understanding of the genetic basis of a disease, allowing for improvements in treatment, detection and prediction of disease susceptibility.

To capture a large amount of genetic variation and increase the chance of detecting associated SNPs, modern GWASs include millions of SNPs sampled from many individuals. GWASs typically examine the independent effect of each SNP on the phenotype by performing a series of statistical hypothesis tests (each SNP requiring the computation of a *test statistic*) under the null hypothesis that no association exists. Rejection of the null hypothesis implies that some form of association exists between a SNP and the phenotype. Including many SNPs in a GWAS gives rise to the *multiple testing problem*, where the cumulative chance of false rejections of the null hypothesis increases with the number of hypothesis tests. GWASs compensate for multiple hypothesis testing by controlling the false positive rate (or the Type 1 error rate), but there is a critical trade-off between suppressing false associations while maximising the number of valid associations.

There are a number of methods for controlling false positives in a multiple testing scenario (4–6) but many of these methods assume the theoretical null distribution of the test statistics which can result in ineffective control of the Type 1 error rate. Permutation testing (PT) is a straightforward and accurate method of minimising the Type 1 error rate but it is very computationally expensive. Although a number of tools (e.g., PLINK (7), GEMMA (8), FaST-LMM (9)) perform GWAS analyses on large data sets, there is a need for a GWAS PT accelerator. FPGAs (Field-Programmable Gate Arrays) are reconfigurable circuits which provide a high level of parallelisation that can be harnessed to accelerate GWAS PT and, although their widespread uptake has been inhibited by their high cost, the availability of cloud-based FPGA platforms providing FPGAs at < US$1.65 per hour (2021) has made powerful FPGAs accessible to bioinformatics labs, and their difficulty to program is now the chief barrier. Freudenthal et al. (10) present GWAS-Flow, a GPU-based accelerator for linear mixed model GWAS PT, but the accelerator is not significantly faster than the more accurate CPU-based implementation GEMMA. Although we have not seen previous work with FPGAs for PT in GWASs, there is work in genome wide association interaction studies (GWAIS). Gundlach et al. (11) present an FPGA-based PT accelerator for GWAIS running on a cluster of 128 low-cost FPGAs. Unfortunately, the purchase and setup of an FPGA cluster is not a viable solution for a typical bioinformatics lab. PBOOST (12) is an example of a GPU-based PT accelerator for GWAIS, but it only supports categorical phenotypes and cannot directly handle confounding variables.

Our paper presents a user-friendly FPGA-based tool to accelerate GWAS permutation testing for continuous phenotypes. An AWS EC2 image is available which can be used to analyse standard PLINK-format files.

## Approach

This work presents an accessible FPGA-based tool for the acceleration of GWAS PT. There is a hardware component (the FPGA design) and a C++ program running on a host computer which manages the FPGA execution using OpenCL (an open-source heterogeneous computing framework) and performs the complex branching logic that is difficult to effectively implement in hardware. The tool accelerates two known PT algorithms: maxT permutation testing (13) and adaptive permutation testing (7, 14). To make the tool accessible to users without access to FPGAs, it is designed to run on an Amazon Web Services (AWS) Elastic Cloud Compute (EC2) FPGA instance. The tool requires PLINK-format as input.

This work uses both a simulated dataset and a real breast cancer dataset to demonstrate the accelerator’s speed and accuracy when compared to PLINK for the two permutation algorithms. The breast cancer dataset consists of 2572 histologically confirmed female African breast cancer cases recruited by the Johannesburg Cancer Study and 1080 matched population-based controls recruited by the AWI-Gen study. Genotyping was done using the Illumina H3Africa Custom Array containing 2.2 million SNPs. Imputation was done through the Sanger Imputation Server using the African Resources Panel. A final dataset of 13.7 million SNPs from 3652 individuals were used for the FPGA accelerator demonstration. As the accelerator is designed to handle continuous phenotypes, the case-control cancer phenotypes were linearised using a generalised linear model.

## Methods

### Association Testing

Linear regression is often used to perform association testing for continuous phenotypes and the *F*-statistic is used to determine the statistical significance of each association test by quantifying the goodness of fit of the linear regression model. For a genotype-phenotype sample of *n* individuals, the *F*-statistic is defined as

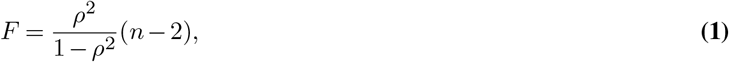

where *ρ* is Pearson’s correlation coefficient

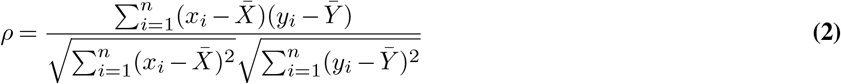

where *Y* is an *n*×1 vector of continuous phenotypes and *X* is an *n*×1 vector of SNP alternate allele counts (*x*_*i*_ ∈ {0, 1, 2, missing}).

### Permutation Testing

Including millions of SNPs in a GWAS increases the chance of detecting associated SNPs, but makes false associations (i.e. Type 1 errors) effectively inevitable when using the traditional *p*-value cut-off *α* = 0.05. Adjusting *α* (the Type 1 error rate) to control for multiple hypothesis testing is therefore critical factor in obtaining valid results from a GWAS.

If the null distribution of the test statistics in a multiple testing scenario is known, more effective control of the Type 1 error rate can be achieved. In practice, however, the true null distribution is unknown. PT controls the Type 1 error rate by simulating the true null distribution of the test statistics using the observed data (14). As PT makes no assumptions about the theoretical null distribution of the test statistics, it can be more effective at controlling the Type 1 error rate than the traditional multiple testing corrections (i.e. Bonferroni, Benjamini and Hochberg).

Randomly permuting phenotypes among the individuals in a GWAS destroys the relationship between genotype and phenotype and represents a sampling under the null hypothesis (1). A PT-adjusted *p*-value is obtained by ranking the observed test statistic among the permuted test statistics. Permuted *p*-values are defined as *p* = *x* + 1*/b* + 1, where *x* is the rank of the observed test statistic and *b* is the number of permutations (15).

Generating PT-adjusted *p*-values with enough resolution to compare against the standard GWAS significance threshold of 5 × 10^*−*8^ (16) needs at least 2 × 10^7^ permutations for each SNP. Consequently, a number of methods have been developed to reduce the computational expense of permutation testing, with two of the more popular methods described here.

#### maxT Permutation Testing

The maxT permutation procedure (13) strongly controls the family-wise error rate (FWER) – the probability of at least one Type 1 error – by generating a null distribution using the *maximum* test statistic from each permutation iteration. A PT-adjusted *p*-value can then be calculated by ranking the observed test statistic against the collection of maximum test statistics. As the maxT method simulates a null distribution using the maximum test statistic of each iteration, much fewer permutations are required to generate a representative null distribution. In fact, at *α* = 0.05, 1000 permutations are typically sufficient, independent of the number of SNPs (17).

#### Adaptive Permutation Testing

Adaptive permutation (7, 14) identifies and stops permuting insignificant SNPs early in the permutation process so that more accurate *p*-values can be calculated for interesting SNPs. Adaptive permutation sets a limit *R* on the number of times the permuted test statistics can exceed the observed statistic. Permuted test statistics are computed until either *R* test statistics are greater than the one observed (requiring *B* permutations) or the maximum number of permutations *b* have been performed. A *p*-value is then estimated as

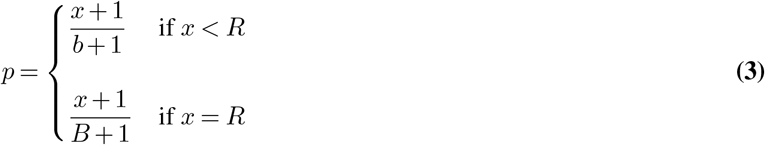

Although the maxT and adaptive permutation algorithms alleviate some of the computational burden of GWAS PT, hardware acceleration can change the scale of the problem that PT can address.

### Architecture of an AWS EC2 FPGA Instance

FPGAs are reconfigurable circuits that can implement arbitrary digital circuits. In contrast to CPUs and GPUs, which have fixed architectures and instruction sets, FPGAs do not have a defined architecture and must be configured to perform a specific function. The adaptability of FPGAs can be exploited to design highly optimised architectures, and FPGAs can be significantly faster and more power efficient than sequentially operating, general-purpose hardware such as CPUs, even though FPGAs are typically clocked an order of magnitude lower. FPGAs are often used as *hardware accelerators* in heterogeneous computing platforms consisting of a *host processor* and one or more hardware accelerators. A heterogeneous architecture allows the host processor to perform complex conditional branching logic (difficult to effectively implement in FPGA hardware), while computationally intensive tasks are offloaded to an optimised hardware accelerator.

AWS EC2 FPGA instances consist of a Xilinx Virtex UltraScale+ VU9P FPGA connected to a host CPU as illustrated by Fig. 1. The host CPU uses OpenCL to communicate with the FPGA via a PCIe interface. OpenCL hides calls to Xilinx runtime (XRT) — a set of communication drivers developed and maintained by Xilinx for Linux. Xilinx provides a pre-configured target platform for AWS FPGA instances which comprises the FPGA logic for the VU9P FPGA together with external DDR memory accessible by both the host processor and the FPGA (i.e. *global memory*). The FPGA logic consists of a static region (which contains I/O, status monitoring and lifecycle management logic) as well as a configurable region for user-developed kernels. The maximum possible frequency of the FPGA design is 250 MHz.

**Fig. 1.**
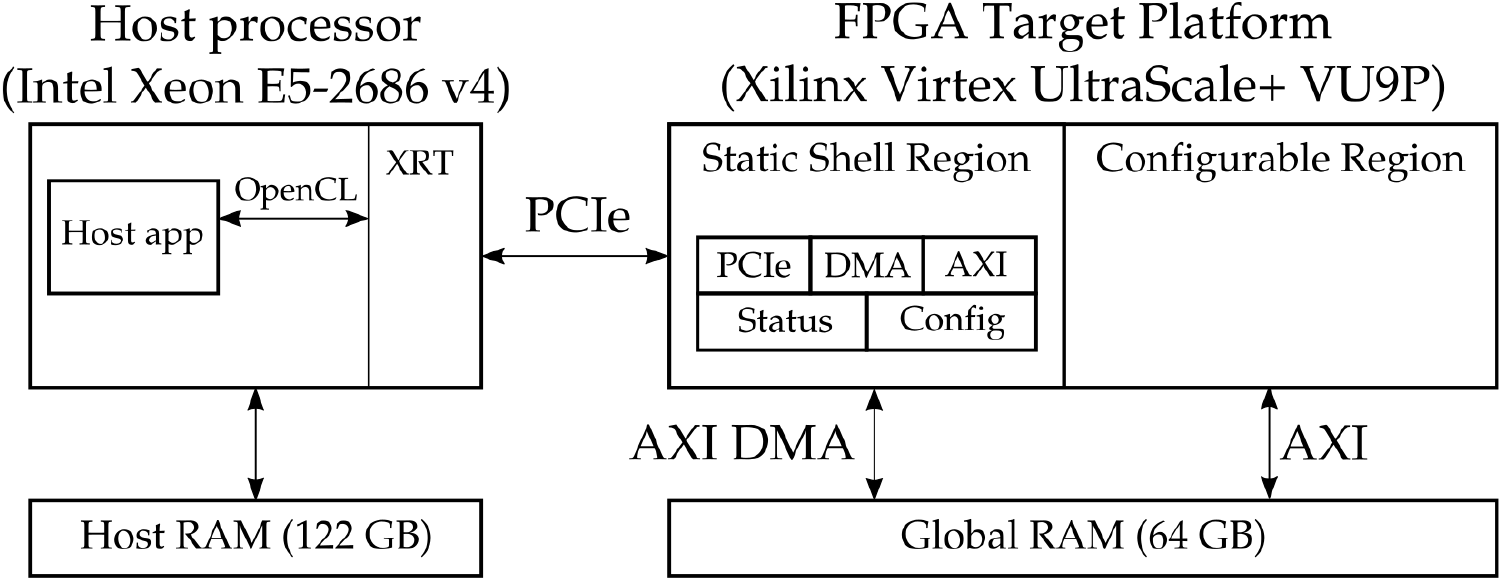
Architecture of an EC2 f1.2xlarge instance.

### FPGA-based PT on a Heterogeneous Platform

PT for one SNP requires the repeated computation of the test statistic *F* with each permutation of the phenotype vector *Y*. When *Y* is permuted, the denominator of *ρ* remains constant while the numerator changes with each permutation. This suggests that the computation of the numerator of *ρ* is the critical part of the permutation testing algorithm. The numerator of *ρ* — the dot product of an *n* × 1 vector of phenotypes (***y***) and an *n* 1× vector of genotypes (***x***) — is defined as:

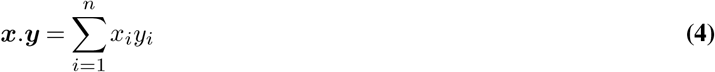

The genotype vector can be expressed as ***x*** = ***a*** − *ā* where, if the additive genotypic model is used, ***a*** is a vector of SNP alternate allele counts for each individual (*a*_*i*_ ∈ {0, 1, 2, missing}). If the GWAS includes *c* covariates for each individual, the phenotype vector can be expressed as ***y*** = ***b −*** *C*(*C*^*T*^ *C*)^*−*1^*C*^*T*^ ***b***, where ***b*** is the real-valued phenotype vector, and *C* is an *n ×* (*c* + 1) matrix of covariates with a prepended column of ones. If covariates are not included, 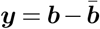

PT for *m* SNPs sampled from *n* individuals requires the computation of the dot product between the permuted phenotype vector ***y***_***p***_ and each genotype vector ***x***_***m***_. The fundamental part of the PT algorithm is a matrix vector multiplication (MVM) between an *m × n* matrix of genotypes and the *n ×* 1 phenotype vector which results in an *m ×* 1 vector of dot products. As MVM is the most computationally expensive part of PT, the most effective way of accelerating PT on a heterogeneous FPGA-CPU platform is to leverage the parallelism of the FPGA to accelerate the MVM, while utilising the CPU to perform the less computationally intensive (although still demanding) tasks such as data preprocessing, management of the FPGA execution and implementation of the PT algorithms.

### FPGA Design

The primary focus of the FPGA design process was the MVM *kernel* that provides an acceptable level of accuracy while minimising latency and maximising throughput to effectively accelerate the MVM calculation. The fact that GWASs aim to identify potentially associated SNPs rather than confirming hypotheses about specific SNPs means than an approximately correct PT *p*-value is typically sufficient for estimating statistical significance. Furthermore, as PT *p*-values are not calculated directly from a test statistic but, rather, from the comparison between test statistics, the computation of extremely high-precision dot products was not required.

The input to the MVM kernel is the genotype data and the phenotype vector. The output is a vector of dot products to be used by the host application to compute the permuted test statistic for each SNP. The genotype data for any reasonably sized GWAS is too large to store in the VU9P FPGA’s onboard RAM (≈48 MB), so it is copied to global memory and streamed to the FPGA for each permutation. The transfer of the genotype data is therefore a major performance bottleneck. To maximise the throughput of the FPGA part of the accelerator, the design: minimises the performance impact of transferring data via global memory by optimising the data types used within the MVM kernel; optimises global memory read/write performance, taking advantage of the inherent parallelism provided by the FPGA hardware; and exploits the architecture of the AWS FPGA instance.

#### Compression of the Genotype Data

The genotype data is much larger than the phenotype vector and the result vector, so the global memory bandwidth is primarily used to transmit the genotype data from host memory to global memory. A major advantage of FPGAs over CPUs and GPUs is their ability to natively operate on arbitrary precision data types. This allows FPGAs to take advantage of the compressed nature of SNP genotype data which can be one of four values (0, 1, 2 or missing). Genotype data can, therefore, be represented with a 2-bit data type, and performing the ***x*** = ***a − ā*** calculation within the MVM kernel (rather than the host application) can significantly reduce the impact of memory bandwidth on the accelerator’s efficiency. As the genotype data does not change throughout permutation testing, ***ā*** is only calculated once for each SNP and, to ensure that the MVM kernel does not have to wait to receive an entire genotype vector before initiating a dot product computation, ***ā*** is calculated by the host application and transferred to the kernel along with the genotype and phenotype data.

#### Fixed-Point Data Types

The MVM kernel uses fixed-point types rather than floating-point types to represent fractional data as fixed-point operations use fewer FPGA hardware resources, have a lower memory footprint and have lower latency than equivalent floating-point operations. Fixed-point designs also consume less power than equivalent floating-point designs.

#### Data Buffering

The phenotype vector is buffered within the kernel’s internal memory so it can be reused for each dot product computation. Not buffering would require the vector to be streamed into the kernel with each SNP vector. This would significantly impact performance as the global memory bandwidth would be split transmitting both the phenotype and genotype data.

A disadvantage is that buffering the phenotype data within the kernel fixes an upper bound on the size of the phenotype buffer when the FPGA design is compiled, limiting the sample size of the datasets that can be analysed. Kernels can be compiled with a large phenotype buffer to accommodate very large sample sizes – on our implementation using the VU9P, the buffer supports samples of ≤262144 individuals. To support larger samples, the host application could be adapted to split the phenotype vector and calculate partial dot products using the MVM kernel (not currently implemented). NB: there is no limit on the number of SNPs.

#### Loop Optimisation

The MVM kernel uses the full width of the global memory interface (512 bits) to reduce the number of global memory accesses during computation, simultaneously increasing kernel parallelism. Loop optimisations allow the MVM kernel to process a 512-bit block of genotype data each clock cycle at the maximum frequency of 250 MHz:

- Loop unrolling supports multiple loop iterations in parallel by replicating the loop hardware several times.
- Loop pipelining is used to overlap loop iterations so a new iteration can be started before the previous iteration completes, allowing the hardware to be maximally utilised.
- Array partitioning is used to allow the phenotype buffer to be accessed in parallel by splitting the array across different RAM resources.

The MVM kernel loops over the rows of the genotype matrix (which store the sampled data of a single SNP) and processes each row in blocks of 512-bits or 256 SNP values. The outer loop, which loops over the rows of the genotype matrix, is not pipelined so as to reduce data dependencies and limit the hardware utilisation of the FPGA design. The middle loop, which loops over a row of the genotype matrix in 512-bit blocks, is pipelined to overlap processing of each 512-bit block. The innermost loop, which performs the multiply-accumulate operations, is fully unrolled to parallelise the processing of each 512-bit block of genotype data.

#### Topological Optimisations

Multiple MVM kernels are instantiated on the FPGA to allow the accelerator to process different blocks of data in parallel. It was found that four MVM kernels maximised the global memory bandwidth, and additional kernels did not result in an increase in performance. The global memory of the AWS FPGA instance is split into four 16 GB banks, so each memory bank is connected to one of the four MVM kernels so as to maximise the system memory bandwidth. All data transferred to a kernel via global memory is transferred using the same memory bank.

### Host Application Design

The C++ host application is optimised to: read the GWAS data from input files; generate the data format expected by the MVM kernels, implement data blocking and buffer management; execute the FPGA kernels with OpenCL; implement the permutation testing algorithms; and record the results.

#### Data Preprocessing

GWAS data is read from industry standard PLINK-formatted files and preprocessed before being sent to the FPGA kernels. A Python 3 script performs covariate/mean adjustment on the phenotype data using the NumPy, pandas and statsmodels packages. The script reads phenotypes from the PLINK *fam* file and writes adjusted phenotypes (either **y** = **b −** *C*(*C*^*T*^ *C*)^*−*1^*C*^*T*^ **b** or 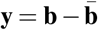, depending on whether a covariate data file is provided) to a file which is read by the host application. The host application converts the phenotypes to the fixed-point format expected by the MVM kernel.

An OpenMP-accelerated (v3.1) algorithm was developed to accelerate genotype preprocessing as it was found to influence the overall run time of the PT process, particularly when the number of SNPs is large and the number of permutations is low (*n*_*p*_ < 1000). The algorithm uses a nested loop pair to process a PLINK *bed* file. The inner loop iterates over each byte of a SNP data block and converts the PLINK-encoded genotype data to the 0, 1, 2 additive genotype format. The inner loop also sums the converted SNP values and the number of non-missing genotypes to calculate the mean of each SNP vector. The outer loop (parallelised with OpenMP) iterates over each SNP in the *bed* file, converts the double-valued mean of each SNP vector to the fixed-point format expected by the MVM kernel and calculates the standard deviation of each SNP vector.

#### Data Blocking

The host application takes advantage of the parallelism of the MVM accelerator by blocking the GWAS dataset so that different blocks of data can be processed concurrently by each of the four MVM kernels. Global memory bandwidth is dependent on the size of the data buffer being transferred, so the host application blocks the genotype data based on the optimal buffer size, which determines the number of kernel invocations as well as the number of dot products computed by each MVM kernel i.e.

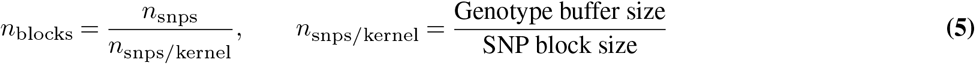

#### Kernel Execution with OpenCL

The host application uses OpenCL (v1.2) API calls to execute the MVM kernels instantiated on the FPGA and to manage data transfer via global memory. OpenCL API calls are used to: initialise the kernels, transfer data to/from the kernels, execute kernel tasks, and synchronise kernel execution based on OpenCL events. All OpenCL API calls used to control kernel execution are asynchronous (i.e. they immediately return after the command has been queued on the OpenCL work queue), so events are used to resolve dependencies between kernel commands. Event-based synchronisation allows the host application to perform other work while OpenCL manages the work queue.

#### Permutation Testing Algorithms

The two PT algorithms were designed with a view towards minimising the overall run time by minimising the idle time of the FPGA kernels. The host application uses multithreading to allow a new permutation to be started while the results of previous permutations are being processed.

The adaptive PT algorithm is more complex than the maxT algorithm as SNPs are periodically removed from the procedure when they do not appear to be significant, which requires the FPGA data buffers to be regenerated. Unlike the maxT algorithm, which has a run time that is affected only by the number of permutations, the adaptive algorithm has a number of parameters, along with the number of permutations, which affect the run time, namely: the SNP dropping interval, the rate at which the dropping interval increases, and the minimum number of permutations for each SNP.

The minimum permutations per SNP (**M-P-S**) is a user-defined parameter. The SNP dropping interval is initially equal to the M-P-S and increases by 20% every 5 drops. At each SNP dropping interval, the host application regenerates the data buffers to remove the SNPs which have been flagged as insignificant and re-runs the data blocking algorithm on the regenerated data. Once enough SNPs have been dropped so that the SNP data can fit into one buffer, the host application transitions to performing one permutation per MVM kernel. During this phase of the algorithm, the host application adaptively sets the size of the genotype data buffers to ensure that the accelerator throughput is maximised. The host application continues to drop SNPs and regenerate the data buffers, but the dropping interval increases more rapidly as only significant SNPs remain and fewer SNPs are removed at each interval.

## Results

The speed and accuracy of the FPGA-accelerated permutation algorithms were compared to PLINK — a popular C/C++ command line tool that implements both maxT and adaptive permutation testing — using two test datasets. PLINK was used as a performance reference as it is one of the most widely used GWAS tools. The FPGA-accelerated permutation algorithms were tested on an AWS EC2 f1.2xlarge instance and PLINK testing was done on a computer cluster with two Intel Xeon Silver 4114 CPUs (each consisting of twenty 2.2 GHz hyper-threaded cores) per node.

The accuracy of the FPGA accelerator was compared to PLINK by comparing the permuted *p*-values to determine whether the same significant SNPs are identified with the same accuracy. As PT is an inherently random process, some variation in the results is expected, but the *p*-values of the most significant SNPs should be very similar. To simulate a realistic GWAS PT scenario, the *p*-value comparison was conducted with maxT *p*-values generated with 1000 permutations and adaptive *p*-values generated with at least 20 million permutations.

These tests were performed without covariates (i.e., the phenotypes were pre-adjusted with covariates) as if a covariate file is included PLINK recalculates the phenotypes for each permutation which significantly reduces the speed of the PT procedure. The FPGA-based algorithms perform the covariate adjustment once at the start of the algorithm, so the inclusion of covariates can be expected to have a minimal effect on the run time of the FPGA-based algorithms.

Two datasets were used for testing and validation of the FPGA-based accelerator: **Dataset 1** — A simulated dataset of 1898052 SNPs sampled from 2018 individuals; **Dataset 2** — A real, imputed dataset of 13742528 SNPs sampled from 3652 individuals.

### maxT Permutation Testing

#### Speed

The run time of the FPGA-based maxT algorithm was compared to that of PLINK running on the cluster. The effect of large-scale parallelisation of PLINK’s maxT implementation was determined by splitting the PLINK maxT algorithm over multiple nodes of the CPU cluster by triggering an instance of PLINK on each of the nodes (where each PLINK instance is required to perform *n*_perms_*/n*_nodes_ maxT permutations) and combining the results (resulting in parallelisation over 40, 200 or 400 CPU cores). The multi-node PLINK tests were only run with the smaller, simulated dataset (Dataset 1), however, as it was not possible to reserve enough time on the cluster to process the real, fully imputed dataset (Dataset 2).

Fig. 2 compares the run time of the FPGA-accelerated maxT algorithm to PLINK’s run time for varying numbers of maxT permutations. The FPGA-based maxT algorithm is at least two orders of magnitude faster than PLINK running on 40 CPU cores for any number of permutations. Furthermore, Fig. 2a demonstrates that the run time of the FPGA-based maxT algorithm is more than an order of magnitude faster than highly parallelised PLINK.

**Fig. 2.**
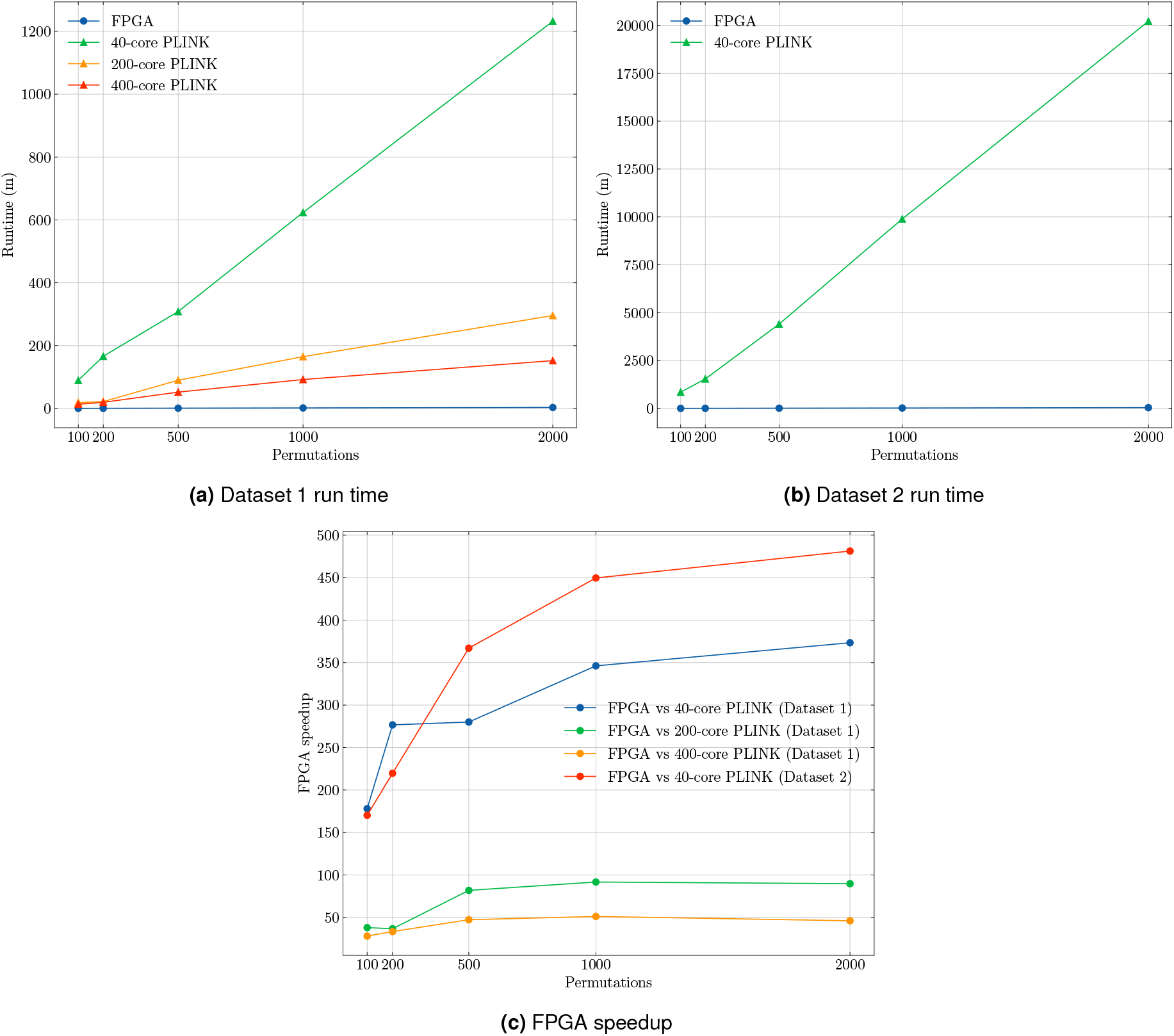
Plots comparing the run time of the FPGA-based maxT algorithm to PLINK.

For 1000 maxT permutations, which provides effective control of the Type 1 error rate for any number of SNPs, the FPGA-based maxT algorithm is 346 times faster than 40-core PLINK for Dataset 1 and 450 times faster than 40-core PLINK for Dataset 2 (see Fig. 2c), suggesting that the FPGA-based maxT accelerator scales well for larger data sets.

#### Accuracy

The accuracy of the FPGA-based maxT algorithm was determined using the significant SNPs identified by PLINK as a reference. Using an arbitrary *p*-value cutoff of *α* = 0.05, PLINK and our tool identified exactly the same 8 significant SNPs from Dataset 1, and 30 significant SNPs from Dataset 2, with no false positives by the FPGA-based maxT algorithm.

### Adaptive Permutation Testing

#### Speed

The data dependencies of the adaptive permutation algorithm prevent the parallelisation of PLINK over multiple computers, so the run time of the FPGA-based adaptive permutation algorithm was compared to the run time of PLINK running on one node of the CPU cluster (40 CPU cores). To determine how the M-P-S affects the run time of the algorithm, Dataset 1 testing was carried out with two M-P-S values (121 and 36) following Che et al. (14), and Dataset 2 with an M-P-S of 121 due to time constraints.

Fig. 3 compares the run time of the FPGA-accelerated adaptive permutation algorithm to PLINK by plotting the respective run times against the maximum permutations per SNP. It can be seen that the FPGA-accelerated algorithm is at least one order of magnitude faster than PLINK for any number of adaptive permutations.

**Fig. 3.**
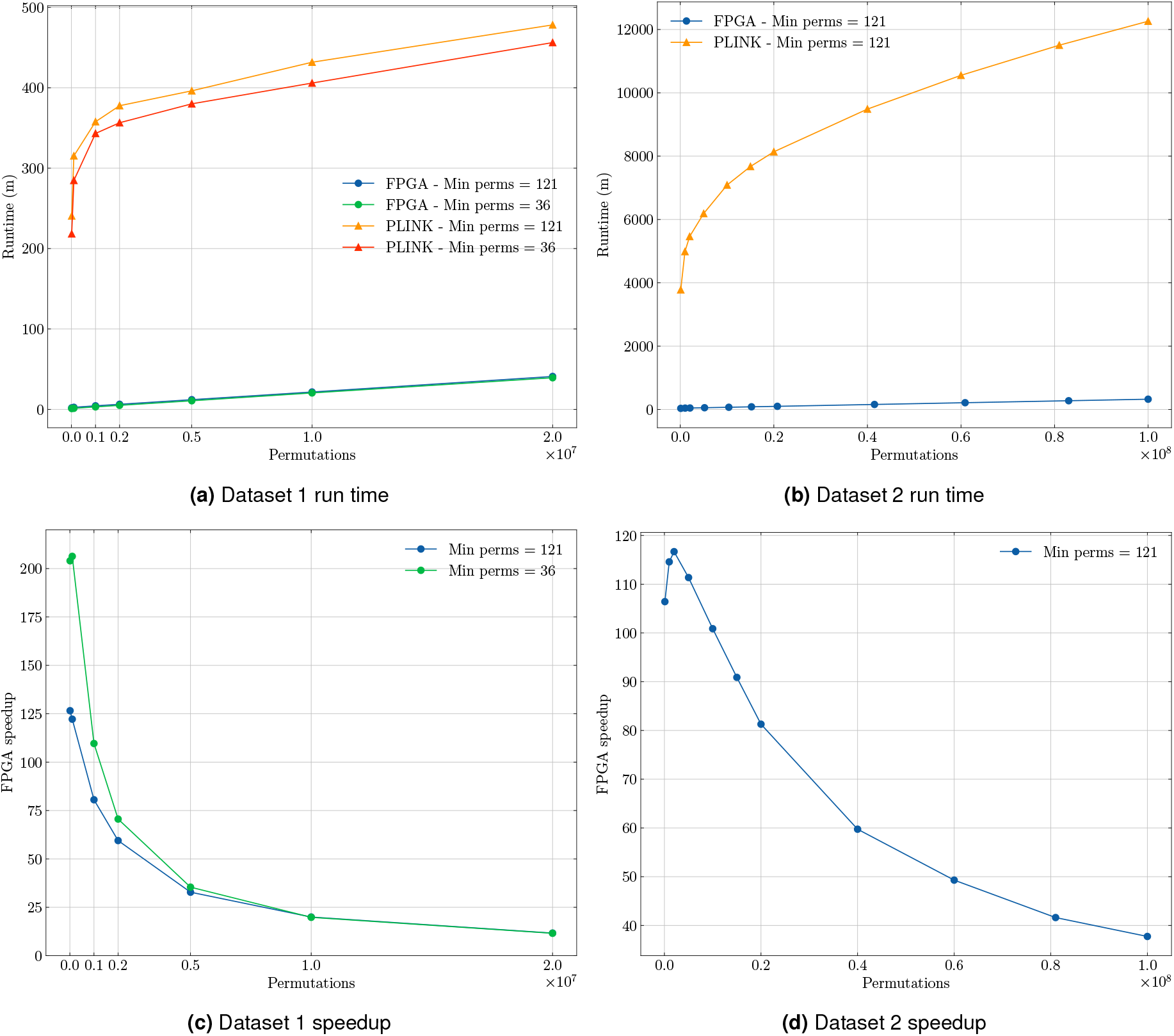
Plots comparing the run time of the FPGA-based adaptive algorithm to PLINK.

Fig. 3 plots the FPGA speedup against the number of permutations for the two test datasets. It is immediately apparent that the speedup achieved for the adaptive permutation algorithm is lower than the speedup achieved for the maxT algorithm. This is caused by two aspects of the adaptive permutation algorithm:

- The algorithm is much more CPU-dependent than the maxT method — the host application must periodically identify and remove insignificant SNPs from the permutation procedure.
- In the later stages of the algorithm, the FPGA accelerator is not operating at peak efficiency due to the use of a suboptimal buffer size.

In spite of this, the FPGA-accelerated adaptive PT algorithm is significantly faster than PLINK for any number of permutations. Fig. 3 shows that the FPGA speedup decreases as the number of adaptive permutations increases – due to a reduction in the efficiency of the FPGA accelerator caused by the use of a sub-optimal buffer size when most of the SNPs have been dropped. Although the FPGA accelerator is not operating optimally in the later stages of the algorithm, it retains a speed advantage over PLINK — particularly in the case of Dataset 2.

For 700 million adaptive permutations of Dataset 2 (an almost unfeasible workload for PLINK running on a 40-core CPU) the run time of the FPGA-accelerated algorithm was 33 hours – significantly less than PLINK’s run time for 100 million permutations of the same dataset (about 8.5 days).

Fig. 3c demonstrates that, although the M-P-S has a significant effect on FPGA speedup when few permutations are performed, this effect becomes negligible as the number of permutations increases. This indicates that the M-P-S only has a significant effect on the run time when many SNPs are undergoing permutation.

#### Accuracy

The accuracy of the FPGA-based adaptive algorithm was determined using the significant SNPs identified by PLINK as a reference. Using an arbitrary *p*-value cutoff of *α* = 1 × 10^*−*7^, PLINK identified 8 significant SNPs after 20 million permutations of Dataset 1, and 50 significant SNPs after 100 million permutations of Dataset 2. The FPGA-based algorithm identified the same significant SNPs as PLINK, although one false positive occurred for Dataset 1. This can be explained by the random nature of the adaptive PT algorithm (an RNG is used to permute the phenotypes) and the fact that more permutations were performed for Dataset 2 so the *p*-values calculated for Dataset 2 are more accurate than those generated for Dataset 1.

## Discussion and Conclusion

This work presents an FPGA-based tool that accelerates PT compared to running parallelised CPU implementations by at least an order of magnitude, implementing both adaptive PT and maxT PT. We provide an AMI that allows user to analyse standard PLINK files – documentation on how to use the AMI is provided at github.com/witseie/fpgaperm. Alternatively, users with a compatible FPGA may download the code and compile themselves.

As the accelerator is designed to run on an AWS EC2 f1.2xlarge instance consisting of a Xilinx Virtex UltraScale+ VU9P FPGA together with an 8-core Intel Xeon host CPU, the PT algorithm was split into a hardware component and a software component. Due to the fact that MVM is the most computationally expensive part of the PT algorithm, the FPGA accelerates the MVM computation, while the host CPU handles preprocessing, managing the FPGA execution and data transfers to/from the FPGA using the OpenCL framework, and implementing the PT algorithms.

PLINK, a popular CPU-based tool that implements both maxT and adaptive permutation testing, was used as a reference with which to compare the speed and accuracy of the FPGA-based algorithms. For a real GWAS dataset of 13.7 million SNPs sampled from 3652 individuals, the run time of PLINK running on 40 Intel Xeon 4114 CPU cores was measured at 164 hours (or approximately 7 days) for 1000 maxT permutations and 204 hours (or around 8.5 days) for 100 million adaptive permutations. In contrast, the run time of the FPGA-based accelerator is 22 minutes for 1000 maxT permutations and 325 minutes for 100 million adaptive permutations resulting in speedups of 447 and 38 respectively. FPGA acceleration enables the handling of workloads that are almost unfeasible for state of the art CPU-based methods. This proves that FPGAs can be effectively employed as bioinformatics accelerators in order to handle the large sample sizes of modern biological datasets thereby opening up new possibilities for bioinformatics research.

While this work has demonstrated that FPGAs can effectively accelerate GWAS permutation testing for continuous phenotypes using simple linear regression, work can be done both to improve the functionality of the accelerator and to improve the performance.

The accelerator cannot currently handle sample sizes of more than 262144 individuals due to the fixed phenotype buffer size. A useful optimisation would be to determine the largest phenotype buffer that does not affect the operating frequency. The accelerator could also be updated to support the analysis of case/control phenotypes which would require the development of a novel FPGA architecture as case/control phenotypes are analysed using logistic regression (or Fisher’s exact test) rather than linear regression.

Further work could be done to improve the efficacy of the FPGA-based adaptive permutation testing algorithm, but this will probably require a redesign of the FPGA part of the accelerator to process multiple permutations in a single invocation of a kernel. This work has not addressed the effect of GPU acceleration on GWAS permutation testing. A comparison between GPU and FPGA-based acceleration of GWAS permutation testing should be undertaken to determine whether comparable speedup can be achieved with GPU acceleration. Additional work could include the analysis of the combined effect of FPGA and GPU acceleration. FPGA acceleration of linear mixed model GWAS permutation testing should be investigated. LMM regression is used to account for genetically related individuals and the fact that LMM regression is significantly slower than simple linear regression means that there is a need for an LMM permutation testing accelerator.

## ACKNOWLEDGEMENTS

We thank Chris Mathew and Michèle Ramsay for making Dataset 2 available: cases from the ERICA-SA Study (https://www.samrc.ac.za/intramural-research-units/evolving-risk-factors-cancers-african-populations-erica-sa) with population-matched controls from the AWI-Gen Study. The ERICA-SA and AWI-Gen studies were approved by the University of the Witwatersrand Human Research Ethics Committee (Medical), protocols 2111154 and 180535.

## FUNDING

YS and SH are partially supported by the Pan-African Bioinformatics Network for H3Africa (National Institutes of Health Common Fund/National Human Genome Research Institute U41HG006941). JTB was supported by H3A AWI-Gen grant (NIH Common Fund/NHGRI, grant #1U54HG006938). MH and WCC were supported by the South African Medical Research Council and the UK Medical Research Council through the UK Government’s Newton Fund, MRC-RFA-SHIP 01-2015. The content is solely the responsibility of the authors and does not necessarily represent the official views of the funders.

## Notes

https://github.com/witseie/fpgaperm

### Competing Interest Statement

The authors have declared no competing interest.

